# The limits of information in precise regulation of early multicellular life cycles

**DOI:** 10.64898/2026.03.19.712848

**Authors:** Hanna Isaksson, William C. Ratcliff, Eric Libby

## Abstract

A key step in the evolution of complex multicellularity is the emergence of regulated life cycles that coordinate growth and reproduction. One potential route toward regulation involves co-opting intrinsic information: cues generated by routine cellular activities such as aging or mechanical stress from growth. Here, we model the simplest form of multicellular organization, linear filaments, to investigate whether intrinsic information can be harnessed to produce regular multicellular life cycles. Based on our analyses, we find that these information sources face an inherent trade-off between flexibility and regularity. Some sources, such as mechanical stress, precisely regulate when reproduction occurs but generate only a single reproductive mode. Others, such as cell age, can in principle produce diverse life cycles but fail to generate any of them reliably. Combining information sources through simple genetic circuits reduces variance in some cases, but the range of achievable life cycles remains constrained. Together these results suggest that while intrinsic information may facilitate early multicellular evolution, there are significant limitations on the degree to which it can be harnessed to evolve tightly-regulated, flexible life cycles. Our work highlights the constraints faced by nascent multicellular organisms and the evolutionary innovations likely required for coordinated multicellular development.

## Introduction

The evolution of multicellular organisms has fundamentally shaped the biosphere by giving rise to novel, increasingly complex forms of life (1, 2). Yet, not all forms of multicellularity are the same: some have hundreds of types of cells arranged in intricate body plans while others are simple clusters of undifferentiated cells (3). Though there may not be a single feature explaining such differences in complexity, a key trait enabling the evolution of complex multicellularity is the ability to tightly coordinate the multicellular life cycle (4, 5). Such coordination typically occurs through transcriptional and translational regulation that brings about reliable patterns of group formation and reproduction. In contrast, simple multicellular organisms often rely on stochastic processes like physical fracturing for group reproduction (6–10), or incidental evolutionary changes like the emergence of non-adhesive cells (11). This raises the question: how do simple multicellular organisms transition from unreliable, stochastic life cycles to tightly regulated development?

One route toward such a transition relies on the incorporation of reliable information into developmental regulation. When available, reliable information provides a significant fitness advantage over stochastic strategies and is therefore expected to be used even when it is costly to sense (12, 13). Examples of this can be found in filamentous cyanobacteria, where nitrogen limitation triggers both heterocyst differentiation and hormogonia formation through coordinated patterns of gene expression and intercellular signaling (14–16). Theoretical studies of multicellular life cycles support this view: regulated life cycles emerge when cells incorporate environmental information, such as population density or external signals, into their reproductive strategies (17–20). In these models, life cycles based on stochastic reproduction are displaced by those that exploit environmental information and connect it to multicellular processes such as group formation or reproduction. A common feature in these studies, however, is the assumption that all cells have equal access to a reliable source of information. This may be problematic as the geometry of even simple multicellular organisms can rapidly create spatial heterogeneity preventing some cells from accessing the environment (3, 6, 21, 22). Another problem with relying on environmental cues is that organisms often live in environments that are relatively constant over the time scale of the life cycle.

A possible alternative way to reliably regulate multicellular life cycles is through the use of intrinsic information, the kind that is generated by the routine activities of cells. Indeed many complex multicellular organisms coordinate development through intricate combinations of gene regulation and molecular diffusion that organize cells into regions with distinct developmental fates (24). Early multicellular developmental programs were probably much simpler, relying on information accessible to individual cells (17–19). For example, in the snowflake yeast model of multicellularity, cells reproduce and remain attached to mother and daughter cells, resulting in a characteristic branched multicellular topology. A consequence of these processes is that older cells are more centrally located (21, 25). With detectable molecular correlates of aging (see, for instance (26, 27)), internal cues could allow cells to infer their age and as a result their position within a multicellular group. Linking this information to the multicellular consequences of cellular behavior (e.g., severing a mother-daughter connection releases a propagule) pro-vides a route through which internal information could be leveraged for the evolution of bespoke multicellular life cycles. Even in relatively consistent environments, life history transitions could be determined by a change in the internal state of the organism (e.g., a size or age-based transition from juvenile to propagule-producing adult (28)). The ability to generate, and act upon, cell-level information in a multicellular developmental context may be an important determinant of whether a nascent multicellular lineage can evolve increasingly complex multicellular phenotypes.

While there are potentially many sources of cell-level information available, their capacity to enact precise life cycles in early multicellularity may be limited. There are likely constraints that would be difficult to overcome without more complex regulation or new sources of information. For instance, a group of 10 cells could theoretically partition into daughter groups in 42 ways, but more complex fragmentation patterns (e.g., splitting into 4 groups of sizes 1, 2, 3, and 4 cells) may be infeasible because they would require multiple, precisely coordinated splitting events. In terms of information, this would mean that multiple cells (at least 3) would have the exact same information at some point in time. Examples of early forms of multicellularity reveal that fragmentation patterns, or reproductive modes, can vary significantly even in the same species experiencing the same selective pressure (see Fig. 1). Theoretical studies have explored the adaptive value of different multicellular reproductive modes (29–32) and found that there are fitness consequences that can select on particular life cycles. Yet, it is unknown to what extent early multicellular groups can actually evolve such precise life cycles to take advantage of these fitness benefits—especially when they must rely on internal, cell-level information.

**Fig. 1.**
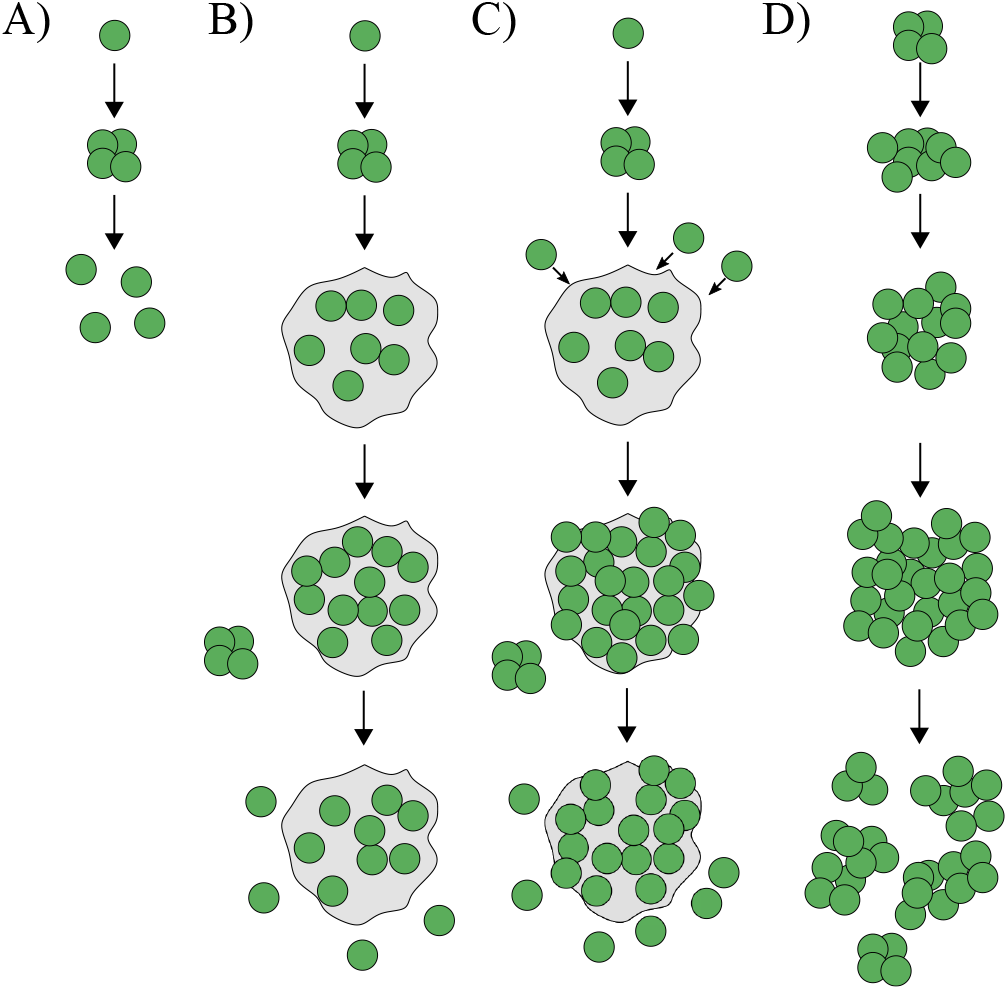
Diversity of evolved multicellular life cycles in a predator-prey experiment. A) A schematic showing a unicellular life cycle in which a single cell grows and then undergoes a rapid series of cell divisions, completely dissociating inti single-celled offspring (propagules). B) A similar life cycle as in (A), but the offspring cells remain embedded in an extracellular matrix (ECM) close to the parent cells, forming a cluster. The cluster reproduces by releasing single-celled propagules. C) A life cycle similar to (B), where cells stay connected via an ECM and reproduce through propagules, but the cluster can also recruit single cells via aggregation. D) A clonal multicellular life cycle that reproduces through fragmentation, maintaining a persistent multicellular stage throughout the life cycle. The diversity among these evolved life cycles highlights the variety that emerges in simple forms of multicellularity, which is a consequence of lacking the ability to regulate the life cycle. The schematic is reconstructed from Herron et. al. 2019 (23).

Here, we consider the simplest form of multicellular group structure, that of one-dimensional multicellular filaments, much like cyanobacteria. We use an agent-based modeling approach to compare the distribution and propagation of information that cells could plausibly have access to, such as cell age and mechanical stress. We examine how these different information sources affect the multicellular life cycles that evolve in terms of their reproducibility and the diversity of evolutionary outcomes. We find that many potential sources of information are not reliable enough to enact a precise life cycle and instead permit a plurality of co-existing life cycles. Ultimately, our results highlight the limitations faced by early multicellular organisms regulating their life cycles via internal cell information.

## Methods

### Simulations of multicellular life cycles

To study the role of information in coordinating multicellular life cycles, we use agent-based simulations of one-dimensional multicellular filaments. We initiate all simulations with a single filament of one cell. Filaments increase in size as cells reproduce via binary fission and stay together post-division. Cell divisions occur independently of one another, and the time until the next reproduction for a given cell is sampled from a normal distribution distribution *N* (1, 0.05) and the time step size for the simulations was 0.01. Filaments fragment according to deterministic rules, such that when cell *i* meets a fragmentation criteria *θ*, the filament breaks at that point. The rules are based on various types of information each with their own criteria *θ*. Specifically, we focus on four types of information accessible to individual cells: cell age, age of cell connection, concentration of a diffusing compound, and mechanical stress. Additionally, we explore fragmentation based on two types of randomness: in the first case, there is a probability of fragmentation each time a cell divides, while in the second case, there is a probability for each cell connection to break at every time step. We end all simulations after 100 fragmentation events.

### Rule-based fragmentation

When a fragmentation event is triggered, which of a cell’s connections is severed depends on the type of information. Below we describe how each information-based fragmentation rule is implemented assuming that cells do not die. For the simulations in which fragmentation occurs via cell death, the cell that satisfies the fragmentation criteria is simply removed and its connections are severed. After each fragmentation event, we select the largest daughter, i.e., the daughter containing the most cells, and follow the growth of that filament.

#### Cell age

For fragmentation based on cell age, an event is triggered when the age of a cell (*a*_*i*_ for cell *i*) exceeds a certain threshold, e.g. *a*_*i*_ > *θ*, which results in the breakage of the oldest connection for the cell *i*, see Fig. 2A. Since assessing the age of cells that grow via binary fission has challenges (33), we use the average age of cell poles as a proxy for cell age, i.e. 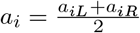, as in (34). We note that we would get similar results using the oldest pole as a measure of aging (e.g. (35)) as they are proportional.

**Fig. 2.**
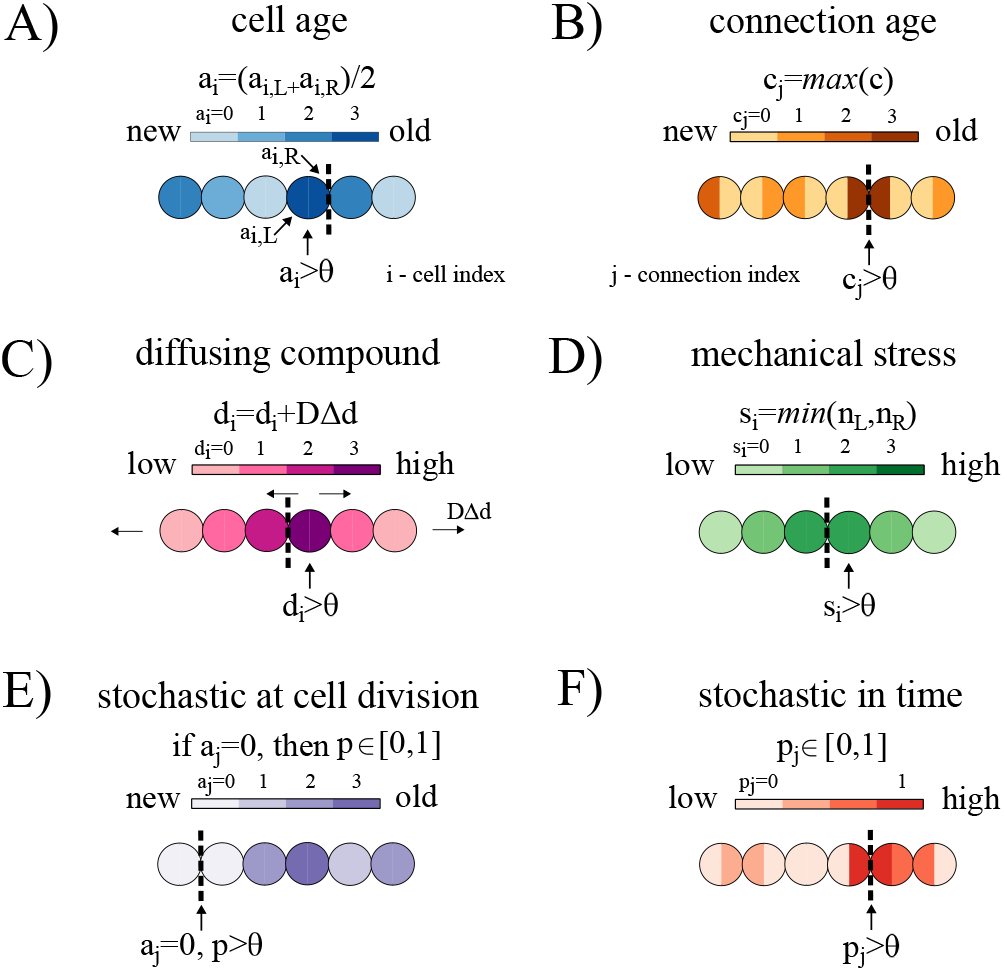
Distribution of internal information in one-dimensional filaments. A) A schematic showing the distribution of cell ages (*a*_*i*_) along a filament, where darker blue indicates older cells. Cell age is defined as the average age of a cell’s poles. B) A schematic showing the distribution of connection ages (*c*_*i*_) along a filament, where darker orange represents older connections. C) A schematic showing the diffusion of a compound (*d*_*i*_) across the filament, where darker pink indicates higher concentration and arrows denote the direction of diffusion (from high to low concentration). D) A schematic showing the distribution of mechanical stress (*s*_*i*_) in a filament, where darker green represents higher strain, which depends on the number of neighboring cells. E) A schematic showing stochastic fragmentation at reproduction, where newly divided cells (light purple) have a probability *p* of breaking their connection to the parent cell, while all other cells have zero probability of fragmentation. F) A schematic showing stochastic fragmentation in time, where each cell connection *j* has a probability *p*_*j*_ of breaking at every time step. Dark red marks connections with a higher fragmentation probability. Each information type is associated with a threshold value *ε*, and fragmentation occurs when this threshold is exceeded.

#### Age of cell connection

Instead of basing fragmentation on cell age, we can base it on the age of connections between cells, see Fig. 2B. For a filament with *n* cells, there are *n* − 1 connections. We let *c*_*j*_ represent the age of connection *j* for *j* = 1, 2,…*n* − 1. The filament breaks whenever a connection is above a certain threshold, i.e. when *c*_*j*_ > *θ* where *θ* is the threshold parameter (see Supplementary information “Cell age vs connection age”).

#### Diffusible compound

For fragmentation based on the diffusion of a compound we let the amount of compound in cell *i* be represented as *d*_*i*_. A fragmentation event is triggered when *d*_*i*_ > *θ* and the filament then breaks on the side of the cell whose neighbor has the highest concentration, see Fig. 2C. The concentration of the compound in a cell increases with the addition of an amount 1 per time step, and the compound also diffuses between cells and from the end cells out to the environment with a diffusion constant *D* = 0.01. Upon cell division, the amount of compound is equally distributed between daughter cells.

#### Mechanical stress

For information concerning the mechanical stress experienced by a cell, we calculate the number of cells downstream on each side of the cell such that *n*_*L*_ is the number on one side and *n*_*R*_ is the number on the other. We note that if a filament has *N* cells then *n*_*L*_ + *n*_*R*_ = *N*− 1. We then define a measure of stress *s*_*i*_ for each cell *i* that is the minimum of *n*_*L*_ and *n*_*R*_, i.e. *s*_*i*_ = min(*n*_*L*_, *n*_*R*_). This biases breaks towards the middle of the filament where stress and torques are likely to be high. If we chose the maximum of *n*_*L*_ and *n*_*R*_ instead of the minimum, it would mean that fragmentation would only occur at terminal cells of the filament. Breakage occurs when *s*_*i*_ > *θ* and we sever the connection with the higher number of cells. For example, if *n*_*L*_ > *n*_*R*_, indicating more cells on the *L* side, then that is the connection that will be severed, see Fig. 2D.

#### Stochastic scenarios

We consider two ways in which filaments may fragment stochastically. First, for the case of stochastic fragmentation at cell division, new connections formed by reproducing cells break with probability *p*, see Fig. 2E. Second, for the case of stochastic fragmentation in time, at each time step of our simulation there is a chance of severing a connection between cells. Specifically, we sample a number *p*_*j*_ for each connection *j* from a uniform distribution *U* (0, 1), and if *p*_*j*_ < *θ* then we break that connection. Thus *θ* represents the probability that a connection will break at a given time step, see Fig. 2F. For simulations in which cell death is studied, we implement cell death by randomly selecting one of the cells surrounding the broken connection for removal.

### Life cycle classification and selection

In order to facilitate comparisons between results obtained using different information types, we select *θ* values for each rule so that the average adult sizes before fragmentation are the same (see Supplementary information “Summary of information types and threshold values” for the *θ* values used in the simulations). When a filament breaks, we classify the observed life cycle based on the number and sizes of offspring. We differentiate between five types of life cycles based on their reproduction mode: 1) unicellular propagules, 2) equal binary split, 3) unequal binary split, 4) complete dissociation, and 5) other. The first type of life cycle, “unicellular propagules”, occurs when fragmentation produces at least one offspring comprised of a single cell and at least one offspring comprised of more than one cell. The second type, “equal binary split”, occurs when there are two offspring filaments, each of which are between 40 − 60% of the size of the parent filament. The third type, “unequal binary split”, occurs when two offspring are produced but they do not satisfy the criterion for “propagule” or “equal binary split”, i.e. the offspring are not of equivalent size. The fourth type, “complete dissociation”, occurs when the entire filament dissociates into single-celled offspring. Finally, fragmentations that do not fall into any of the previous four types are categorized as “other” (see Supplementary information “Classification of reproduction modes” for more details).

## Results

To determine the extent to which early multicellular life cycles can be tuned, we developed an agent-based model of a simple filamentous life cycle (see Methods “Simulation of multicellular life cycles”). Cells within filaments reproduce via binary fission, causing filaments to grow exponentially. Filaments reproduce whenever a connection between two cells breaks. In our modeling framework, such fragmentation events are triggered by specific rules that represent simple forms of developmental regulation. These rules rely on information that cells reproducing via binary fission might plausibly access. For this study, we consider four types of such information: 1. cell age, 2. cell connection age, 3. the amount of a diffusing compound, and 4. mechanical stress. Figure 2 illustrates how fragmentation is determined by each type of information (see also Methods: “Rule-based fragmentation”). As a benchmark for these deterministic rules, we also consider two stochastic fragmentation scenarios in which events occur randomly at the moment of cell reproduction or randomly over time, independent of reproduction. This modeling framework allows us to evaluate the extent to which cell-level information can regulate the structure of a multicellular life cycle.

### Regulated adult size

The life cycles of one-dimensional filaments have two main traits that might be regulated in response to selection: the size at which a filament reproduces (adult size) and the distribution of offspring sizes (reproduction mode, see Fig. 3A). We first examine the capacity of information-based rules to regulate the adult size of filaments, and later consider the reproductive mode. We set a target adult size of 32 cells and found the parameter (*θ*) for each fragmentation rule that minimized the average squared error. Figure 3B shows the resulting adult size distribution for each rule (see Supplementary information “Summary of information types and threshold values” for the parameter settings). For rules based on connection age and mechanical stress, fragmentation can be precisely tuned to occur at 32 cells with little to no error (for mechanical stress *σ* = 0, for connection age *σ* < 0.35). In contrast, the other rules produce life cycles that rarely reproduce at exactly 32 cells (< 3% for cell age and < 5% for diffusing compound), with adult sizes ranging from as few as 2 cells to over 100 cells. Variation is even higher under the two stochastic scenarios, which produce similar adult size distributions (*p* < 1*e* − 9 using a Brown–Forsythe test when grouping stochastic scenarios and comparing with deterministic rules). The qualitative patterns among the different rules persist across other choices of target adult size (see Supplementary information “Adult size distribution for varying *θ*”).

**Fig. 3.**
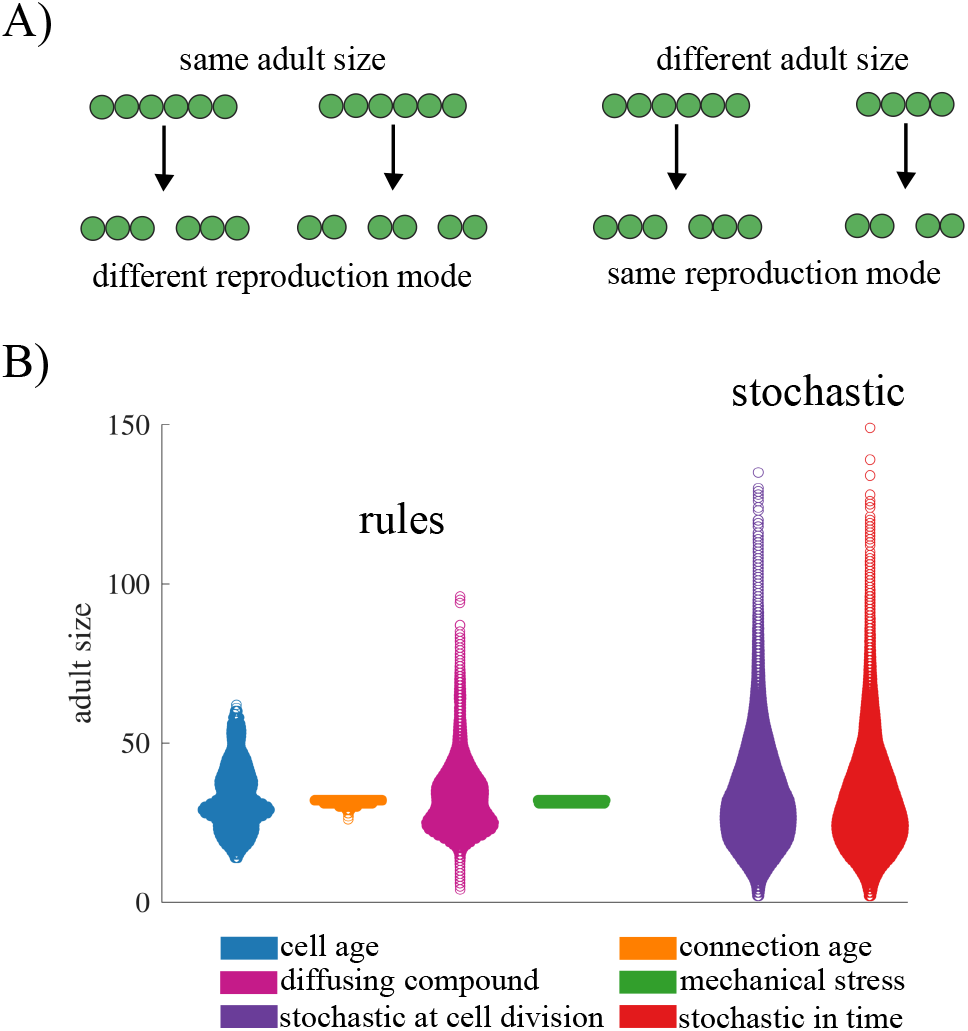
The effects of information types on adult size distributions. A) A schematic illustrating two aspects of a filamentous life cycle that can be regulated: the adult size at fragmentation or the number of offspring per fragmentation event. B) A swarm plot showing the distribution of adult sizes for fragmentation rules based on different types of internal information, as well as for stochastic fragmentation. In all cases, the threshold *θ* was adjusted so that the mean adult size was approximately 32. Rules based on mechanical stress and connection age generate highly regulated life cycles that reproduce consistently at the same adult size. In contrast, rules based on cell age or a diffusible compound produce greater variability in adult size. Largest variation in adult size was observed for stochastic fragmentation.

### Regulated fragmentation mode

Next, we explore the effects of different information types on the ability to generate a specific reproduction mode, so that reproduction of the life cycle follows a consistent pattern. Although there are many ways for a filament to fragment (30), we focus on four main reproduction modes: unicellular propagules, binary split, unequal binary split, and complete dissociation (see Figure 4A and Methods “Life cycle classification and selection”). We also include a category of “other” to include modes of reproduction that do not fall into one of the four categories. Using the parameter values selected for an average adult size of 32 cells, we characterize how often we observe each of the five categories of reproduction modes from 100,000 fragmentation events (see Figure 4B).

**Fig. 4.**
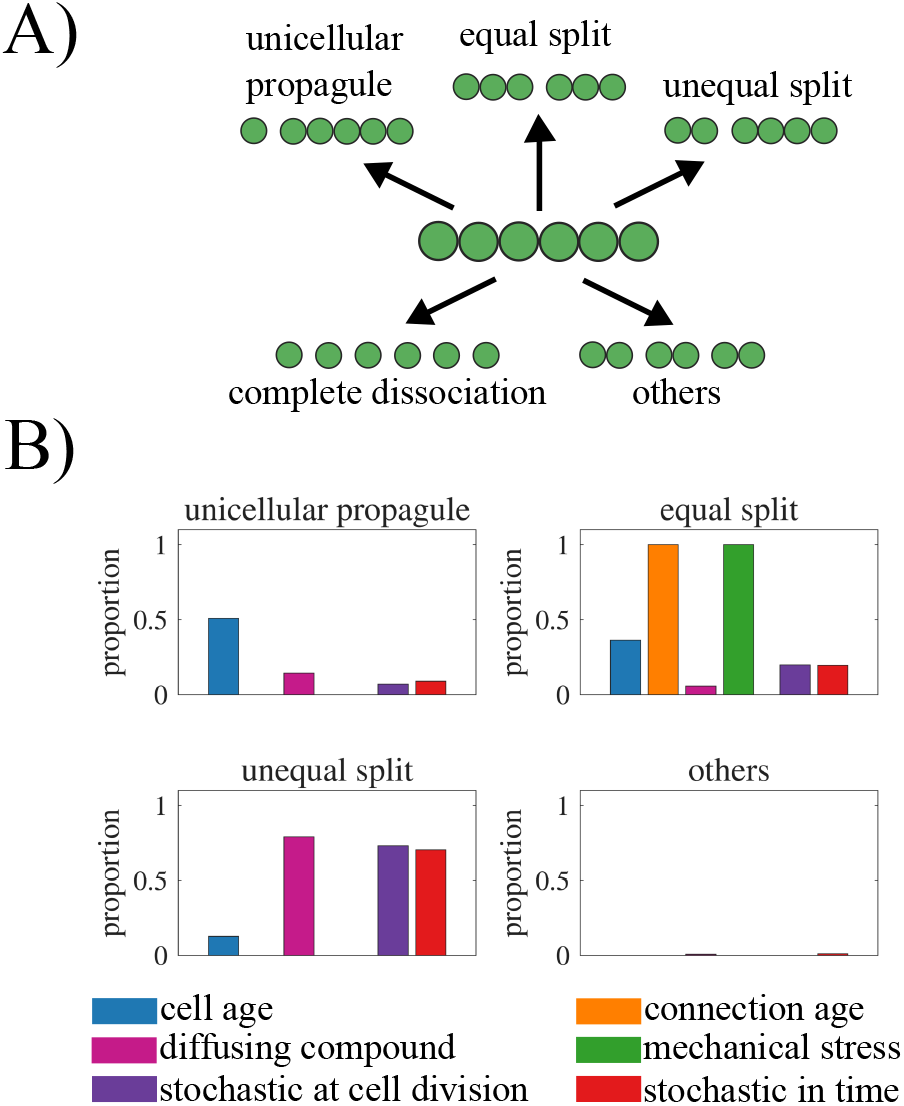
Internal information and reproduction modes. A) A schematic illustrating the classification of life cycles according to reproduction mode: (1) unicellular propagule, (2) equal binary split, (3) unequal binary split, (4) complete dissociation, and (5) other. B) A bar graph showing the frequency of each reproduction mode observed across 10,000 fragmentation events, with colors indicating the corresponding information type. Complete dissociation is not shown, as it was not observed for any information type. The figure shows that the type of internal information governing fragmentation determines the distribution of reproduction modes, highlighting how different information rules give rise to distinct life cycle patterns.

Figure 4B shows that we find instances of all categories of reproduction modes across the various fragmentation rules except complete dissociation, which never occurred in our simulations. In order for a filament to reproduce via complete dissociation all connections between cells need to be severed simultaneously. For this to happen, all cells would need to have the same information simultaneously, but none of the information sources we consider develops in that manner. Of the remaining reproductive modes, only the “other” category includes instances of multiple cells simultaneously severing a connection, leading to more than two offspring filaments. We observe such reproductive events only when fragmentation is based on a diffusing compound (7.2% of fragmentations) or when it is stochastic in time (10.4% of the fragmentations).

Based on the distributions of the observed reproduction modes, we identify a couple of patterns. First, fragmentations based on cell connection age or mechanical stress only give rise to life cycles that reproduce via equal split (binary fission). Since mechanical stress in our model is always highest for the cell in the middle of the filament, a rule that associates fragmentation with stress will result in equal splits. For cell connection age, the oldest connection also tends to be in the middle of the filament due to the symmetry of cell divisions— but sometimes it may be higher elsewhere (34). Depending on the target adult size, the oldest connection rule can produce life cycles that reproduce via unequal split, though equal split is still more common (see Supplementary information “Reproduction modes for varying *θ*”).

Second, the remaining rules reproduce by all modes (except complete dissociation) but they have different biases. For example, the majority of fragmentations according to cell age occur via unicellular propagules (51%) while the rule incorporating a diffusing compound most often leads to unequal split (79%). The two types of stochastic fragmentation act similarly to each other, favoring unequal split followed by equal split and then single cell propagules. We note that these biases are similar when we select for filaments with different target adult sizes (see Supplementary information “Reproduction modes for varying *θ*”) with the exception of the diffusible compound rule. As the target adult size increases the rule reproduces less often via equal splits and more often via single cell propagules and unequal splits.

### Interaction between adult size and reproduction mode

In the previous analyses we selected fragmentation rules for a specific average adult size (32 cells). With the exceptions of rules using mechanical stress or cell connection age, we observed a plurality of different life cycles with the vast majority of reproductive events occurring in filaments that were not of the selected adult size. It could be that reproductions at these different sizes may follow more predictable patterns, thus we consider how the distribution of reproduction modes depends on the actual size at which filaments reproduce. For each rule, we bin filament reproductions based on the adult size and track how often each reproductive mode is realized. Figure 5 shows that the distributions change with the size at reproduction for all rules, except mechanical stress and oldest connection.

**Fig. 5.**
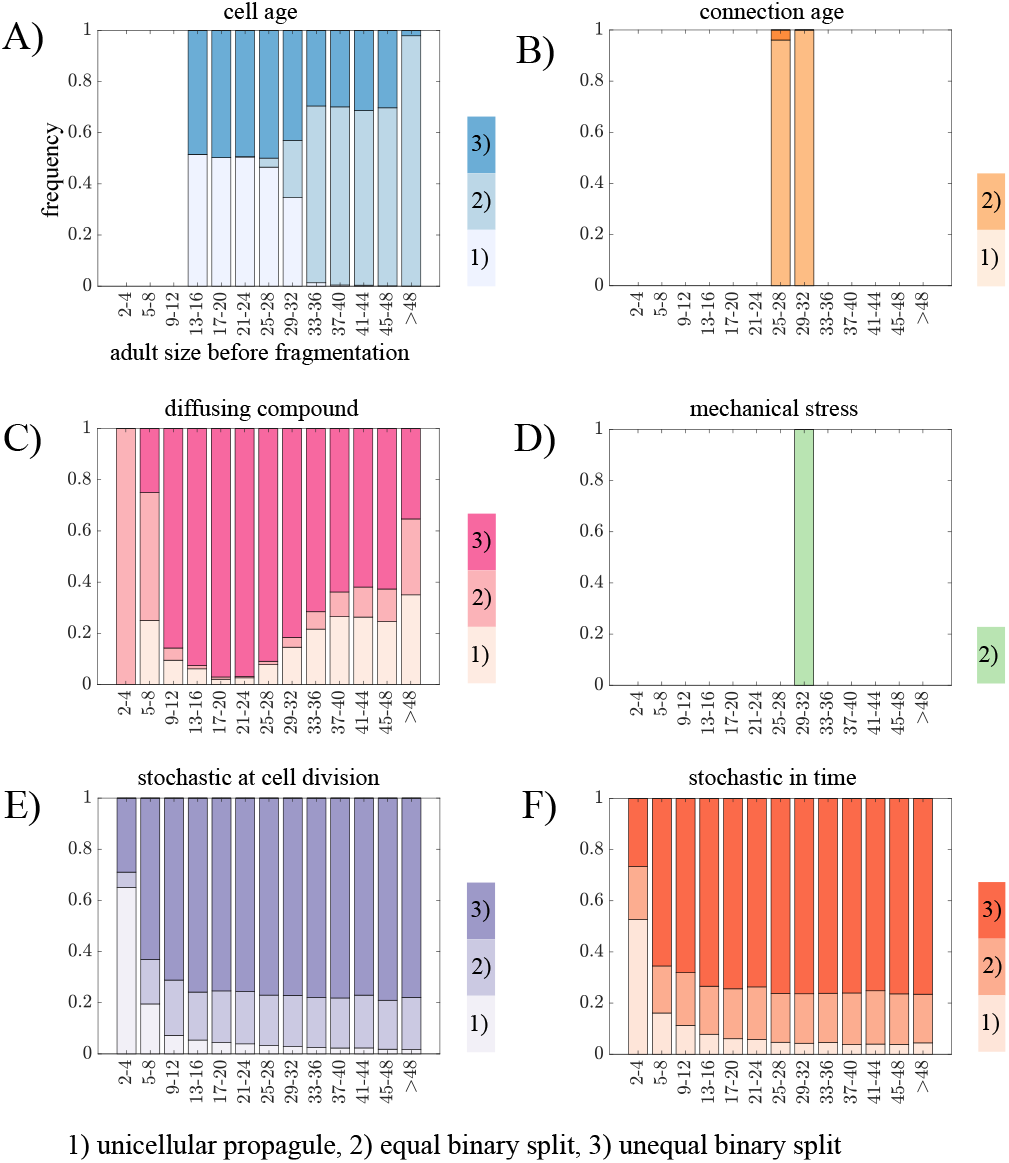
Variations in reproduction modes for different adult sizes. A-F) Bar graphs showing the fraction of fragmentation events resulting in each reproduction mode — (1) unicellular propagule, (2) equal binary split, and (3) unequal binary split — when mother filaments are grouped into adult-size classes. Complete dissociation was never observed, and only reproduction modes occurring with a frequency greater than 1% are shown. Panels correspond to different fragmentation rules:(A) cell age, (B) connection age, (C) diffusing compound, (D) mechanical stress, stochastic fragmentation at cell division, and (F) stochastic fragmentation at each time step. The figure shows that most information types produce substantial variation in both adult size and reproduction mode, indicating that fragmentation dynamics — and consequently life cycle structure — depend strongly on the size of the parent filament.

We observe a general pattern that appears in the rule based on cell age as well as the stochastic fragmentation scenarios. In small filaments a common mode of reproduction is via unicellular propagules, while in larger filaments reproduction occurs more often via equal split (in the cell age rule) and unequal split (in the two stochastic fragmentation scenarios). For the filaments that fragment according to the cell age rule, this pattern emerges as a natural consequence of cells in filaments reproducing repeatedly via binary fission: the oldest cells are the terminal cells and the next oldest cells are close to the middle of the filament (34). So once a single cell propagule has been released (because it was the oldest) it sets the stage for the next fragmentation to be an equal split. For the stochastic rules the pattern emerges as a statistical consequence of the number of cells and connections: the only way to produce a single cell propagule is to sever a connection with a terminal cell and the probability of this type of event decreases as filaments increase in size. Similarly, as filaments increase in size there are more connections that can be severed to produce an unequal split.

The fragmentation rule based on a diffusing compound does not follow any of the previously observed patterns, i.e. 1. only equal binary splits or 2. single cell propagules for small filaments and binary split for large filaments. Instead, this rule creates a large fraction of unequal binary splits for most adult sizes, with an increased tendency to produce single cell propagules for long filaments. The reason for this is that initially, the cells in the middle of the filament have the highest concentration of the diffusing compound. Upon fragmentation, the new terminal cell (previously a middle cell) will have a high concentration of the compound. However, since the compound diffuses from areas of high levels of compound to low, terminal cells gradually lose the compound both outward into the environment and inward toward the filament’s center. As a result, the highest compound level is no longer at the terminal cell, instead it is somewhere between the terminal cell and the center, resulting in an unequal binary split.

### The effects of death and aging on fragmentation

Until now, we have considered fragmentation events that sever connections between cells in the absence of cell death. This potentially allows the cell that triggered the fragmentation event to remain in the largest filament and influence future fragmentations. Here we consider what changes occur when the fragmentation event results in permanent removal of the cell (see Methods “Rule-based fragmentation” for details on implementation in the various fragmentation rules). The results of our simulations show that while including cell death does not significantly affect the adult size distribution (see Supplementary information “Adult size distribution for fragmentation with death”), it does change the observed reproduction modes—except for the rule based on mechanical stress (see Figure 6).

**Fig. 6.**
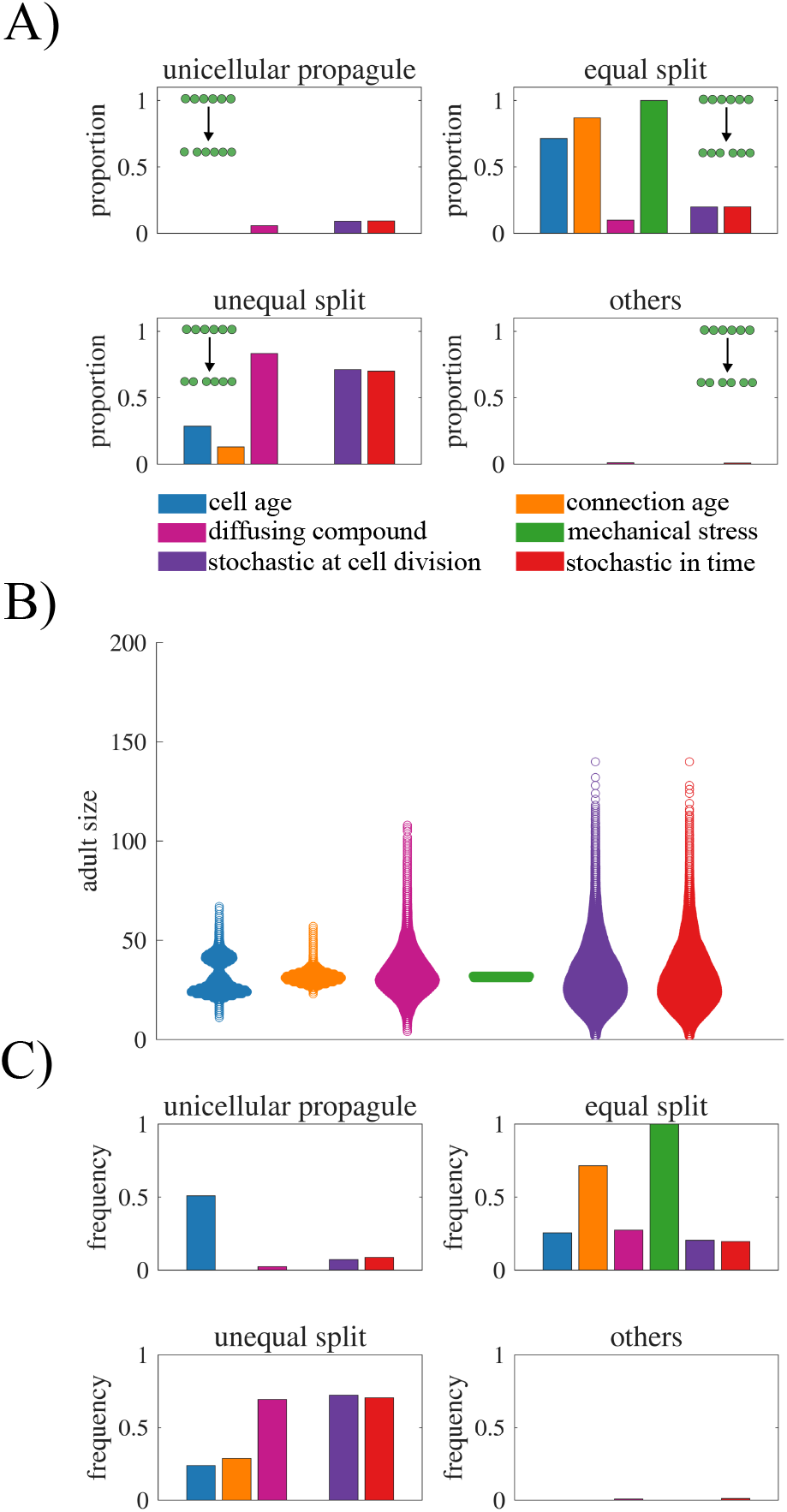
Effects cell death and aging. A) Bar graphs showing the frequency of each life cycle observed for different information types when fragmentation occurs as a result of cell death. A notable decrease in unicellular propagule frequency is observed for age-related information types. B) Scatter plot showing the adult size distribution when older cells have slower growth rate. C) Bar graphs showing the reproduction modes resulting from age-dependent growth. The figure illustrates that cell death constrains which reproduction modes can occur, whereas cell aging introduces additional variability, reducing regulation in the filamentous life cycle.

A prominent effect of incorporating cell death is a reduction of single cell propagules. This effect is particularly evident in the fragmentation rule based on cell age. In the absence of cell death, single cell propagules were the most common reproductive mode. When cell death is introduced, single cell propagules are not observed, and instead, equal splits emerges as the dominant mode of reproduction. This shift is a consequence of the age distribution within a filament. The oldest cells typically are the end cells and so removing them (via cell death) results in essentially the same filament but with one less cell. While technically the filament fragments, one daughter is a dead cell, and so it does not represent a viable filament offspring. Following this event, the filament will continue to grow until it fragments again—this time from an older cell closer to the middle.

Besides cell death, another natural process that might affect the distribution of information is cell aging/senescence, and in particular its effects on the rate of cell reproduction. Past work has shown that age-induced slowing of cell reproduction can affect the rate of adaptation in multicellular filaments (34). To explore how it might shape information and life cycle regulation, we implemented a model in which older cells divide more slowly. We do this by adding the cell’s current age to the expected reproduction time, delaying the next division event for older cells. We find that incorporating cell aging introduced stochasticity into the system increasing the variability in adult sizes. We also found differences in the frequency of reproduction modes in rules using connection age and diffusing compound. Without aging all reproductions using connection age information resulted in equal binary splits, but with aging 27.75% of fragmentation events resulted in unequal binary splits. The diffusing compound rule showed the opposite trend, shifting toward more equal binary divisions, with equal splits increasing from 5.76% to 52.78%, and unequal splits decreasing from 79.14% to 46.06%.

### Combining information types

In our simulations, we have thus far considered fragmentation events triggered by a rule based on a single type of information. Here, we study the consequences of using rules that combine two types of information. To integrate two types of rules, we draw inspiration from systems biology where connections are made between boolean logic and genetic regulation (36–40). Thus, we consider two qualitatively different ways to combine information types: AND and OR logic (see Fig. 7A). In the case of AND logic, fragmentation occurs only when both rules are satisfied. In OR logic, fragmentation occurs when either rule is satisfied. As a case study, we analyze the consequences of these AND/OR rules combining cell age and the amount of a diffusing compound, though additional combinations are explored in the SI (see Supplementary Information, “Combined information – All combinations”).

**Fig. 7.**
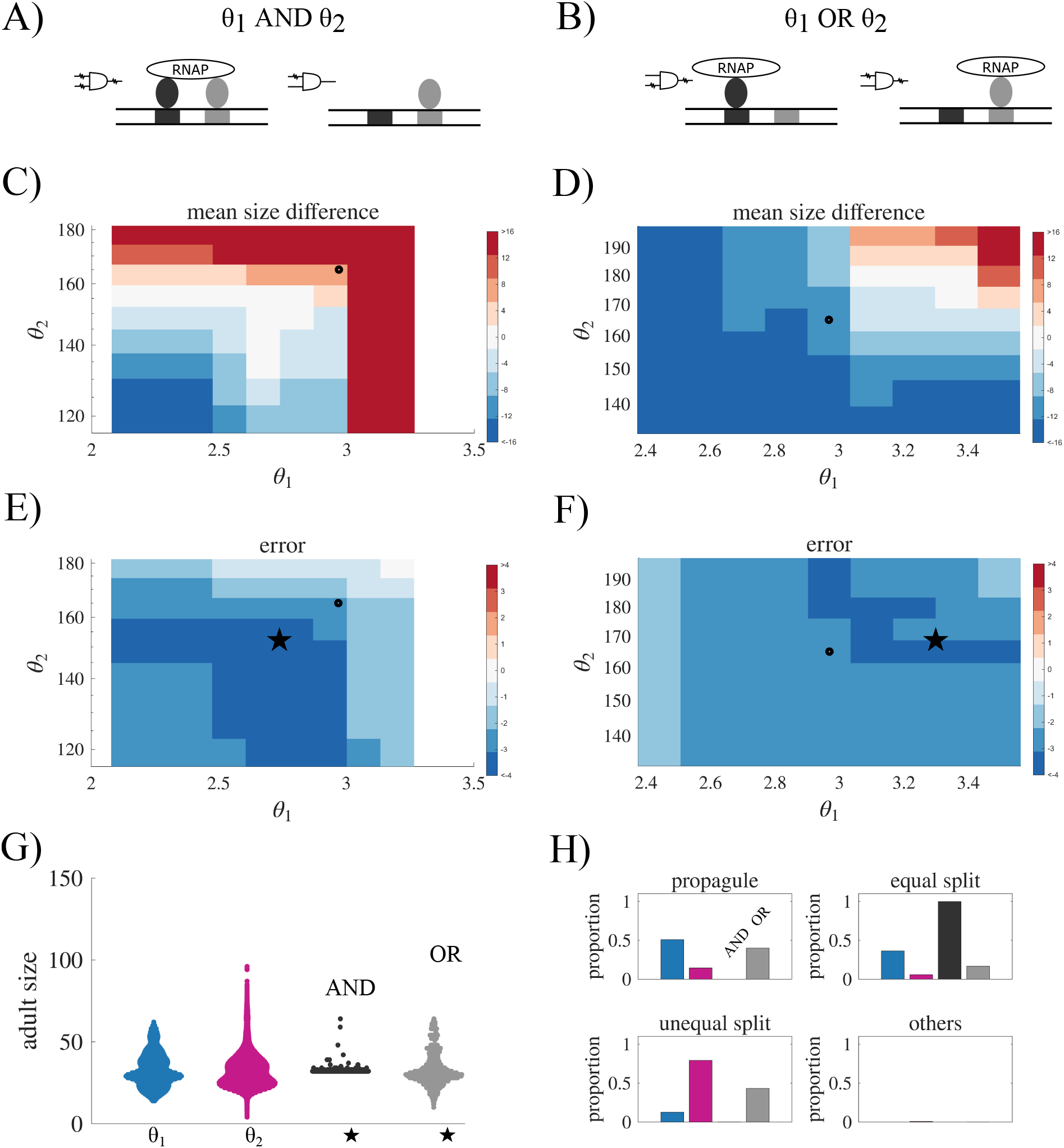
Combined information types: cell age and diffusible compound. A) A schematic illustrating two information types combined into a fragmentation rule using a logical AND, where both conditions must be met. B) A schematic illustrating the OR combination, where either condition is sufficient. C) Contour plot showing the deviation in mean adult size across combinations of thresholds *σ*_1_ (cell age) and *σ*_2_ (diffusible compound) using the AND function. The black circle marks the optimized thresholds that yield an adult size of 32 when each information type is used in isolation. D) Similar to C, but for the OR combination. E) Contour plot showing adult size variation (L2 norm) for the AND combination. The black circle is the same as in C, and the black star indicates a set of threshold values (*σ*_1_, *σ*_2_) that generate low error. F) Similar to E, but for the OR combination. G) Scatter plot showing the adult size distribution for each information type used in isolation and combined at the thresholds indicated by stars in panels E and F. H) Bar graph showing the reproduction modes for each information type in isolation and for the combined thresholds. The figure shows that AND combinations tend to overshoot the target adult size for a large set of threshold values, whereas OR combinations tend to produce too small adults. Overall, combining information types can improve life cycle regulation compared to using either information type alone by reducing adult size variation and increasing the frequency of certain reproduction modes.

We scan a range of thresholds for cell age (*θ*_1_) and the amount of a diffusing compound (*θ*_2_). For each combination we simulate 100,000 fragmentations and compute the error from a target adult size of 32 cells. Fig. 7B) shows the difference between the average adult size at fragmentation and the target adult size. The black dot indicates the optimal threshold parameters when each source of information is used in isolation. In the majority of cases the incorporation of another source of information, causes the average adult size to deviate from the target size. If we compare the AND rule versus the OR rule, we see that the AND rule more often causes the average adult size to increase above the target size (colored red) while the OR rule causes it to decrease (colored blue). This occurs because for the AND rule to activate both conditions must be met for fragmentation to occur, making the rule more restrictive. By comparison, the OR rule is more permissive because fragmentation can occur if either condition is satisfied.

Although both AND/OR rules cause deviations in the average adult size, they reduce the overall error (as calculated by the mean squared error). Figs. 7E) and F) show that the error is less than the best single source of information over the entire parameter space of (*θ*_1_, *θ*_2_) combinations that we evaluated. The reason the error is reduced is because combining information sources reduces the variance in the distribution of adult sizes (see Fig. 7G), and this reduced variation lowers the overall error.

Another consequence of incorporating more information types is that it can alter the pattern of reproductive modes. Fig 7H shows that both the AND and OR rule deviate from the patterns produced by each single source of information. For example, the OR function produces a fragmentation profile that resembles an average of the profiles from the individual information types. This may be expected, as the OR condition is satisfied whenever either threshold is met, meaning that each fragmentation event can effectively be attributed to one of the two individual rules. In contrast, the AND function leads to a significant increase in the frequency of equal binary splits, a reproductive mode that was rarely observed when each information source is used in isolation. Together these results indicate that combining fragmentation rules can lead to new life cycles that have more regular patterns in terms of adult size and reproduction mode (see Fig. 7E).

## Discussion

A key step in the evolution of complex multicellularity is the establishment of regulated life cycles. Underlying this transition is a shift in the drivers of multicellular reproduction, from random events largely dictated by extrinsic factors to more precise mechanisms that incorporate cell-level cues such as age, mechanical stress, or the concentration of diffusible compounds. Here, we examined the capacity of simple multicellularity, in the form of multicellular filaments, to regulate their life cycles using the types of intrinsic information cells might plausibly access. We simulated thousands of multicellular generations in which fragmentation followed deterministic rules based on cell-level information. Our results revealed fundamental challenges inherent with information sources in terms of reliability and flexibility. Information sources either failed to precisely regulate the size at which filaments reproduce or were able to exactly regulate the size but could only produce a single type of life cycle in terms of the mode of multicellular reproduction. Combining sources of information led to life cycles with less variation in terms of adult sizes but with limitations in the types of life cycles that could be produced. These results highlight the challenges faced by early multicellularity in terms of the structure and precision of their life cycles and suggest that the evolution of regulated life cycles likely required novel mechanisms for developmental coordination and communication among cells.

The space of possible multicellular life cycles is vast, corresponding to different combinations of reproductive size and cell allocation. In our analyses, we focused on groups of 32 cells which could theoretically partition into daughter groups in 8348 different ways, yet we found that only a few of these could actually be realized and fewer still could be regularly produced. Consider a life cycle that reproduces via complete dissociation, in which the group reaches a characteristic adult size and dissociates into single-cell offspring. This requires all cells to possess the same information, which does not occur without some synchronous signal that can be sensed by all cells, such as a significant change in the environment. The only life cycle we found that could be reliably produced uses binary fission for group reproduction, where the group splits in half at a specific adult size. This life cycle could be generated by information concerning mechanical stress or the age of cell-cell connections. Because these sources of informa-tion distinguish the center of a filament from its ends, they could not produce other life cycle structures with different reproductive modes, such as multiple offspring, single cell propagules, or unequal splits. While other types of information could produce these reproductive modes, they could not produce any of them exclusively: in these cases, we observed a plurality of reproductive modes and sizes at which fragmentation occurred. Together, these results highlight that the space of multicellular life cycles capable of being regularly produced by co-opting cell-level information may be extremely limited (see Figure 8).

**Fig. 8.**
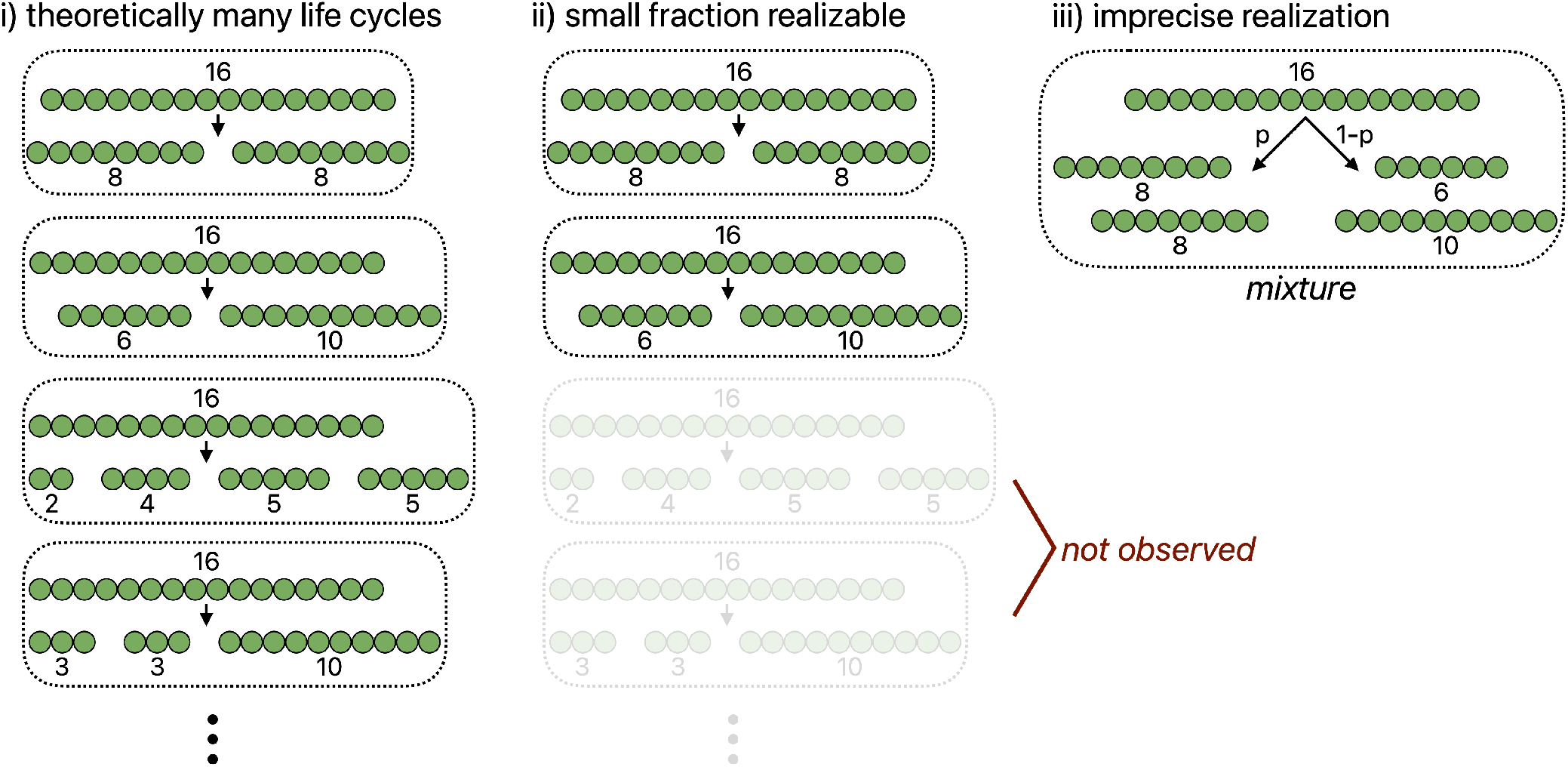
Challenges in producing precise multicellular life cycles. A schematic summarizes the implications of our results concerning the extent to which early multicellular life cycles can be precisely regulated. First, there is the combinatorial space of life cycles that are possible. For illustrative purposes, panel i) depicts filaments that reproduce upon reaching 16 cells, although other adult sizes are also possible (e.g., reproduction at 15 or 17 cells). Despite this large space of possible life cycles, only a small subset can be produced when reproduction is determined without global coordination, i.e., by random events or by rules based on cell-level information. Some life cycles, particularly those with multiple births, are difficult to achieve because they would require different cells to have identical sources of information (the “not observed” in panel ii)). Finally, even among the life cycles that can be realized, precise implementation is often challenging, resulting in mixtures of life cycles. In the schematic example in panel iii), a proportion *p* of fragmentation events follows one mode, while the remaining 1 − *p* follow another.

Natural life history processes such as aging and cell death could potentially be coopted by multicellular organisms to coordinate life cycles, and we used our model to assess this possibility. In certain multicellular life cycles, fragmentation occurs through cell death. Using our models we observed that the realized life cycles varied depending on whether the fragmenting cell died or survived. The primary difference was the loss of life cycles that reproduced via single-cell propagules, which is a common mode of reproduction in complex multicellular organisms (41). This life cycle was particularly impacted because it often results from information in a terminal cell surpassing a threshold: if that cell dies, no viable offspring is produced, and the filament merely sheds an end cell rather than reproducing. In contrast, the effects of aging were less pronounced. The main result was a shift in the distribution of reproductive modes in simulations where fragmentation is based on connection age or diffusing compounds. Contrary to expectation, life cycles based on cell age were less affected when aging led to slower reproductive times. One reason is that both the rule and the slowing of reproduction have a similar effect of biasing fragmentation towards the oldest cells. In both cases of cell death and aging, we again observed strong constraints on the control and tuning of multicellular life cycles.

Although some sources of information were unable to regulate the multicellular life cycle in isolation, combining information sources yielded improvements. For example, cell age and the concentration of a diffusing compound could be combined via simple AND logic to produce a more regular life cycle, one whose reproductive mode was infrequently observed when either source was used independently. Notably, this combination mimicked the fragmentation patterns that resulted from mechanical stress, suggesting convergent constraints on achievable life cycle structures. Thus, even when information sources are combined, strong limitations persist on which multicellular life cycles can be realized.

For our analyses, we studied an extremely simple multicellular life cycle in which cells are arranged in linear filaments and reproduction is determined by severing a single connection. Our intent was to study a “best-case” scenario: multicellular groups with permanent cell bonds and deterministic geometries ensure that information is heritable across generations. Because groups follow the same developmental patterns, they recapitulate the same spatial and temporal structure, allowing cell-level information to reliably correlate with position within the group. For instance, a cell of a given age will consistently find itself at the center or ends of the filament. This predictability breaks down in aggregative multicellularity, where cells can form and break bonds with multiple neighbors, disrupting the positional correlates of information like aging. Additionally, in aggregative organisms, multiple bonds may need to be severed to cause group fragmentation rather than a single one as in a filament, and the number of bonds may vary stochastically depending on which cells happen to be attached to one another. Similarly, the extra dimensions of multicellular groups growing in sheets or snowflake-like branching structures may pose further challenges for cell-level information to reliably indicate group size or facilitate particular reproductive modes. Beyond geometric considerations, cell-level traits can also alter how information manifests. In the snowflake yeast model of multicellularity, for instance, cells reproduce via budding instead of binary fission, and this difference has been shown to impact the distribution of cell ages within filaments(21) as well as broader evolutionary outcomes such as the rate of adaptation (34). Consequently, for early multicellular organisms, a group’s initial physical arrangement and its cell-level life history traits will interact to shape both information generation and its utilization in facilitating multicellular life cycles.

In this paper, we uncovered limits to the regulation of early multicellular life cycles, revealing a fundamental barrier that multicellular evolution must overcome. The information that cells can glean from their own growth and reproduction, though sufficient to produce a narrow range of life cycles, cannot support the diversity of reproductive modes and developmental patterns observed across multicellular life. Animal development illustrates how this limitation has been circumvented: dedicated cell-cell signaling pathways such as Notch, Wnt, and Hedgehog provide information that no single cell could generate on its own, while morphogen gradients establish positional information across tissues through specialized ligand-receptor systems that have been tuned by selection to reliably convey spatial coordinates (42, 43). These mechanisms are integrated through complex gene regulatory networks that buffer noise and coordinate developmental timing across the organism (44, 45). None of this machinery is a simple extension of unicellular sensory capacity. Rather, it represents the evolution of bespoke developmental systems operating at the multicellular scale. The constraints we identify may thus represent an early bottleneck in the evolution of complex multicellularity, one that can only be overcome through the origin of dedicated machinery for generating, transmitting, and interpreting information among cells.

## Acknowledgments

H.I. and E.L. acknowledge support from the Swedish Research Council 2022-04124. W.C.R. acknowledges support for this work from NIH grant R35GM138030.

## Bibliography

1. John Maynard Smith and Eors Szathmary. The major transitions in evolution. OUP Oxford, 1997.

2. Matthew D Herron, Peter L Conlin, and William C Ratcliff. The evolution of multicellularity. CRC Press, 2022.

3. Andrew H Knoll. The multiple origins of complex multicellularity. Annual Review of Earth and Planetary Sciences, 39:217–239, 2011.

4. Jordi Van Gestel and Corina E Tarnita. On the origin of biological construction, with a focus on multicellularity. Proceedings of the National Academy of Sciences, 114(42):11018–11026, 2017.

5. Jordi van Gestel, Martin Ackermann, and Andreas Wagner. Microbial life cycles link global modularity in regulation to mosaic evolution. Nature Ecology & Evolution, 3(8):1184–1196, 2019.

6. Shane Jacobeen, Jennifer T Pentz, Elyes C Graba, Colin G Brandys, William C Ratcliff, and Peter J Yunker. Cellular packing, mechanical stress and the evolution of multicellularity. Nature physics, 14(3):286–290, 2018.

7. Piyush Nanda, Julien Barrere, Thomas LaBar, and Andrew W Murray. A dynamic network model predicts the phenotypes of multicellular clusters from cellular properties. Current Biology, 34(12):2672–2683, 2024.

8. Raymond C Highsmith. Reproduction by fragmentation in corals. Marine ecology progress series, 7:207–226, 1982.

9. Janie L Wulff. Asexual fragmentation, genotype success, and population dynamics of erect branching sponges. Journal of Experimental Marine Biology and Ecology, 149(2):227–247, 1991.

10. Antonia Herrero, Joel Stavans, and Enrique Flores. The multicellular nature of filamentous heterocyst-forming cyanobacteria. FEMS microbiology reviews, 40(6):831–854, 2016.

11. Paul B Rainey and Benjamin Kerr. Cheats as first propagules: a new hypothesis for the evolution of individuality during the transition from single cells to multicellularity. BioEssays, 32(10):872–880, 2010.

12. Matina C Donaldson-Matasci, Carl T Bergstrom, and Michael Lachmann. The fitness value of information. Oikos, 119(2):219–230, 2010. doi: 10.1111/j.1600-0706.2009.17781.x.

13. Edo Kussell and Stanislas Leibler. Phenotypic diversity, population growth, and information in fluctuating environments. Science, 309(5743):2075–2078, 2005. doi: 10.1126/science.1114383.

14. Enrique Flores and Antonia Herrero. Compartmentalized function through cell differentiation in filamentous cyanobacteria. Nature Reviews Microbiology, 8(1):39–50, 2010.

15. Cheng-Cai Zhang, Sophie Laurent, Samer Sakr, Ling Peng, and Sylvie Bédu. Heterocyst differentiation and pattern formation in cyanobacteria: a chorus of signals. Molecular microbiology, 59(2):367–375, 2006.

16. John C Meeks and Jeff Elhai. Regulation of cellular differentiation in filamentous cyanobacteria in free-living and plant-associated symbiotic growth states. Microbiology and Molecular Biology Reviews, 66(1):94–121, 2002. doi: 10.1128/MMBR.66.1.94-121.2002.

17. Eric Libby and Paul B Rainey. Eco-evolutionary feedback and the tuning of proto-developmental life cycles. PLoS One, 8(12):e82274, 2013.

18. Emma Wolinsky and Eric Libby. Evolution of regulated phenotypic expression during a transition to multicellularity. Evolutionary Ecology, 30:235–250, 2016.

19. Enrico Sandro Colizzi, Renske MA Vroomans, and Roeland MH Merks. Evolution of multi-cellularity by collective integration of spatial information. Elife, 9:e56349, 2020.

20. Merlijn Staps, Jordi Van Gestel, and Corina E Tarnita. Emergence of diverse life cycles and life histories at the origin of multicellularity. Nature ecology & evolution, 3(8):1197–1205, 2019.

21. Eric Libby, William Ratcliff, Michael Travisano, and Ben Kerr. Geometry shapes evolution of early multicellularity. PLoS computational biology, 10(9):e1003803, 2014.

22. Salva Duran-Nebreda and Ricard Solé. Emergence of multicellularity in a model of cell growth, death and aggregation under size-dependent selection. Journal of the Royal Society Interface, 12(102):20140982, 2015.

23. Matthew D Herron, Joshua M Borin, Jacob C Boswell, Jillian Walker, I-Chen Kimberly Chen, Charles A Knox, Margrethe Boyd, Frank Rosenzweig, and William C Ratcliff. De novo origins of multicellularity in response to predation. Scientific reports, 9(1):2328, 2019.

24. Julian Lewis. From signals to patterns: space, time, and mathematics in developmental biology. Science, 322(5900):399–403, 2008.

25. William C Ratcliff, Johnathon D Fankhauser, David W Rogers, Duncan Greig, and Michael Travisano. Origins of multicellular evolvability in snowflake yeast. Nature communications, 6(1):6102, 2015.

26. Georges E Janssens and Liesbeth M Veenhoff. Evidence for the hallmarks of human aging in replicatively aging yeast. Microbial Cell, 3(7):263, 2016.

27. Steffen Fehrmann, Camille Paoletti, Youlian Goulev, Andrei Ungureanu, Hugo Aguilaniu, and Gilles Charvin. Aging yeast cells undergo a sharp entry into senescence unrelated to the loss of mitochondrial membrane potential. Cell reports, 5(6):1589–1599, 2013.

28. Stephen C Stearns. The evolution of life histories. Oxford university press, 1998.

29. Corina E Tarnita, Clifford H Taubes, and Martin A Nowak. Evolutionary construction by staying together and coming together. Journal of theoretical biology, 320:10–22, 2013.

30. Yuriy Pichugin, Jorge Peña, Paul B Rainey, and Arne Traulsen. Fragmentation modes and the evolution of life cycles. PLoS computational biology, 13(11):e1005860, 2017.

31. William C Ratcliff, Matthew Herron, Peter L Conlin, and Eric Libby. Nascent life cycles and the emergence of higher-level individuality. Philosophical Transactions of the Royal Society B: Biological Sciences, 372(1735):20160420, 2017.

32. Hanna Isaksson, Åke Brännström, and Eric Libby. Minor variations in multicellular life cycles have major effects on adaptation. PLoS Computational Biology, 19(4):e1010698, 2023.

33. Eric J Stewart, Richard Madden, Gregory Paul, and François Taddei. Aging and death in an organism that reproduces by morphologically symmetric division. PLoS Biol, 3(2):e45, 2005.

34. Hanna Isaksson, Peter L Conlin, Ben Kerr, William C Ratcliff, and Eric Libby. The consequences of budding versus binary fission on adaptation and aging in primitive multicellularity. Genes, 12(5):661, 2021.

35. Søren Vedel, Harry Nunns, Andrej Košmrlj, Szabolcs Semsey, and Ala Trusina. Asymmetric damage segregation constitutes an emergent population-level stress response. Cell systems, 3(2):187–198, 2016.

36. Nicolas E Buchler, Ulrich Gerland, and Terence Hwa. On schemes of combinatorial transcription logic. Proceedings of the National Academy of Sciences, 100(9):5136–5141, 2003.

37. Eric Libby, Theodore J Perkins, and Peter S Swain. Noisy information processing through transcriptional regulation. Proceedings of the National Academy of Sciences, 104(17): 7151–7156, 2007.

38. Sacha AFT van Hijum, Marnix H Medema, and Oscar P Kuipers. Mechanisms and evolution of control logic in prokaryotic transcriptional regulation. Microbiology and Molecular Biology Reviews, 73(3):481–509, 2009.

39. Stuart Kauffman, Carsten Peterson, Björn Samuelsson, and Carl Troein. Random boolean network models and the yeast transcriptional network. Proceedings of the National Academy of Sciences, 100(25):14796–14799, 2003.

40. Timothy S Gardner, Charles R Cantor, and James J Collins. Construction of a genetic toggle switch in escherichia coli. Nature, 403(6767):339–342, 2000.

41. Thibaut Brunet and Nicole King. The origin of animal multicellularity and cell differentiation. Developmental cell, 43(2):124–140, 2017.

42. Eric Dessaud, Andrew P McMahon, and James Briscoe. Pattern formation in the vertebrate neural tube: a sonic hedgehog morphogen-regulated transcriptional network. Development, 135(15):2489–2503, 2008. doi: 10.1242/dev.009324.

43. James Briscoe and Stephen Small. Morphogen rules: design principles of gradient-mediated embryo patterning. Development, 142(23):3996–4009, 2015. doi: 10.1242/dev.129452.

44. Eric H Davidson, Jonathan P Rast, Paola Oliveri, Andrew Ransick, Cristina Calestani, Chiou-Hwa Yuh, Takuya Minokawa, Gabriele Amore, Veronica Hinman, Cesar Arenas-Mena, et al. A genomic regulatory network for development. Science, 295(5560):1669– 1678, 2002. doi: 10.1126/science.1069883.

45. Nikolaos Balaskas, Ana Ribeiro, Jasmina Panovska, Eric Dessaud, Noriaki Sasai, Karen M Page, James Briscoe, and Vanessa Ribes. Gene regulatory logic for reading the Sonic Hedgehog signaling gradient in the vertebrate neural tube. Cell, 148(1-2):273–284, 2012. doi: 10.1016/j.cell.2011.10.047.

