## Supplementary material for "The limits of information in precise regulation of early multicellular life cycles": SI

### Supplementary information

**Cell age vs connection age.** In our model, each cell is represented by two poles — left and right — and we track the age of these poles. As cells reproduce via binary fission, newly formed daughter cells inherit one old pole from the mother cell and gets one newly formed pole at the side where the new cell wall is created. This asymmetric inheritance leads to a structured distribution of pole ages across the filament (34). We define cell age as the average of the left and right pole ages for each cell. In contrast, connection age refers to the age of the connection between two neighboring cells, which is equal to any of the age of the adjacent poles. Thus, any connection between two cells inherits the same age as the closest pole. To illustrate the difference between these two age-based rules, we provide a schematic diagram (Fig. S1) showing the development of filaments growing via binary fission.

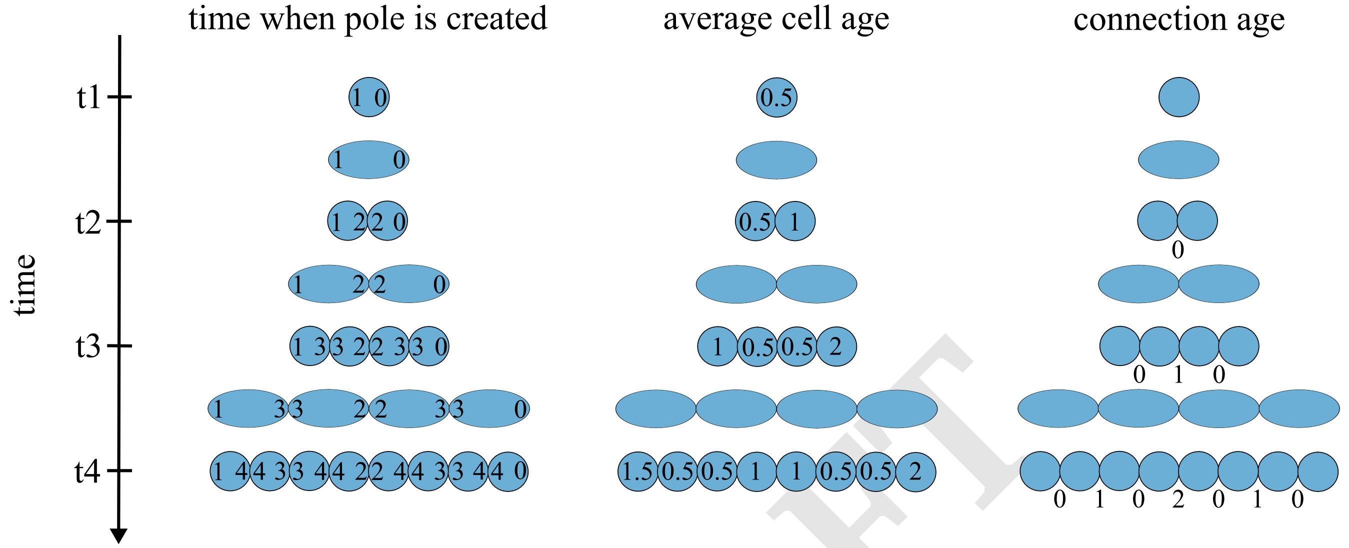


**Fig. S1. Pole age, cell age, and connection age in a filament undergoing binary fission.** This diagram illustrates the distinction between aging at the level of individual cells versus intercellular connections. Left: A filament of cells with the time of creation indicated for each pole. Middle: Cell age, calculated as the average of the left and right pole ages. Right: Connection age, assigned as the age of any of the adjacent poles.

DRAFT

**Summary of information types and threshold values.** We study the effects of six different types of internal cell information on filament fragmentation. The first four information types are deterministic, while the last two are stochastic. For each information type, the fragmentation threshold $\theta$ is calibrated to produce an average adult filament size of 32 cells in the simulations. Once a cell's internal value reaches this threshold, the filament fragments at that cell.

In a one-dimensional filament, each internal cell has two neighbors (except the terminal cells), and fragmentation requires selecting which connection to sever. For the age-based rules, fragmentation occurs on the side of the cell that has the oldest pole. In the case of a diffusing compound, the filament breaks on the side where the concentration of the compound is highest. When fragmentation is governed by mechanical stress, we break the filament on the side where there are more neighboring cells, assuming that a heavier load on that side makes it more prone to breaking.

For the two stochastic information types, the rules differ. In the stochastic-at-reproduction case, fragmentation always occurs between the two most recently produced daughter cells. In contrast, for the stochastic-in-time rule, when a cell reaches the threshold, one of its two connections is chosen at random for breakage.

The parameters and values used for each information type are summarized in Tab. S1.

**Table S1. Information types and threshold values used in the simulations.** Summary of the six internal information types used to determine filament fragmentation. For type we defined by how the fragmentation is implemented and the calibrated threshold $\theta$ that triggers fragmentation. Thresholds are set such that the average adult filament size is 32 cells in all cases.

| Information type | Implementation | Parameter | Threshold ($\theta$) |
|---|---|---|---|
| Cell age | Oldest pole | $a_i$ | 2.9688 |
| Connection age | Oldest pole | $c_i$ | 4.25 |
| Diffusing compound | Highest concentration | $d_i$ | 165 |
| Mechanical stress | Highest number of neighbors | $s_i$ | 14 |
| Stochastic at cell division | The new cell connection | $a_j$ and $p$ | 0.1156 |
| Stochastic in time | Random pole | $p_j$ | 8.8672e-4 |

**Classification of reproduction modes.** At each fragmentation event, the resulting reproduction mode has to be classified. We distinguish between the four reproduction modes: "unicellular propagules", "equal binary split", "unequal binary split", and "complete dissociation". In particular, the difference between equal and unequal binary split is that for equal binary split, the size of each of the two offspring is not smaller than $40\%$ of the adult size and not larger than $60\%$ of the adult size. Reproduction modes that do not fall into these four categories are classified as "other", which could, for example, be three offspring of different sizes all of which are larger than one. The procedure for classifying the result from a fragmentation event is shown in Fig. S2.

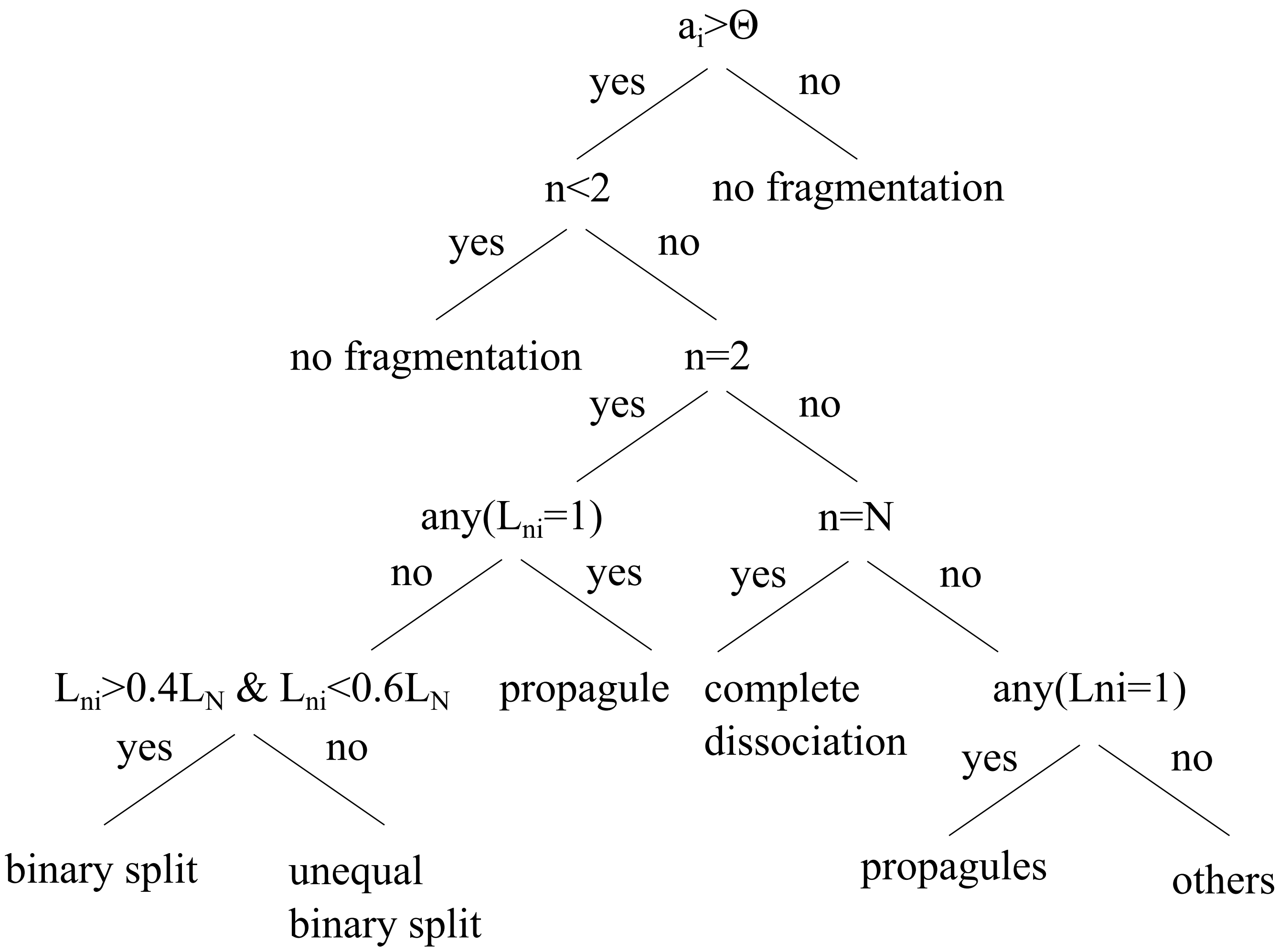


**Fig. S2. Decision tree for classifying reproduction modes.** Upon each fragmentation event the resulting reproduction mode has to be classified. The classification is done according to the decision tree shown in this figure. A fragmentation event happens if the condition $\theta$ is fulfilled and if the number of resulting offspring is more than one. Here, $n$ is the number of offspring, $N$ is the adult size, and $L_{ni}$ is the size of offspring $i$.

**Adult size distribution for varying $\theta$.** To assess the robustness of our selected threshold values, we provide the distribution of resulting adult filament sizes for each information type across three different threshold values, see Fig. S3. As expected, increasing the threshold generally leads to larger adult sizes. However, the overall adult size pattern is preserved across thresholds. Fragmentation based on mechanical stress remains completely deterministic, consistently producing a single adult size. Connection age leads to low variability, whereas cell age and the diffusing compound result in broader size distributions. The two stochastic information types — stochastic at cell division and stochastic in time — are similar to one another and exhibit the highest variation in adult size across all threshold settings.

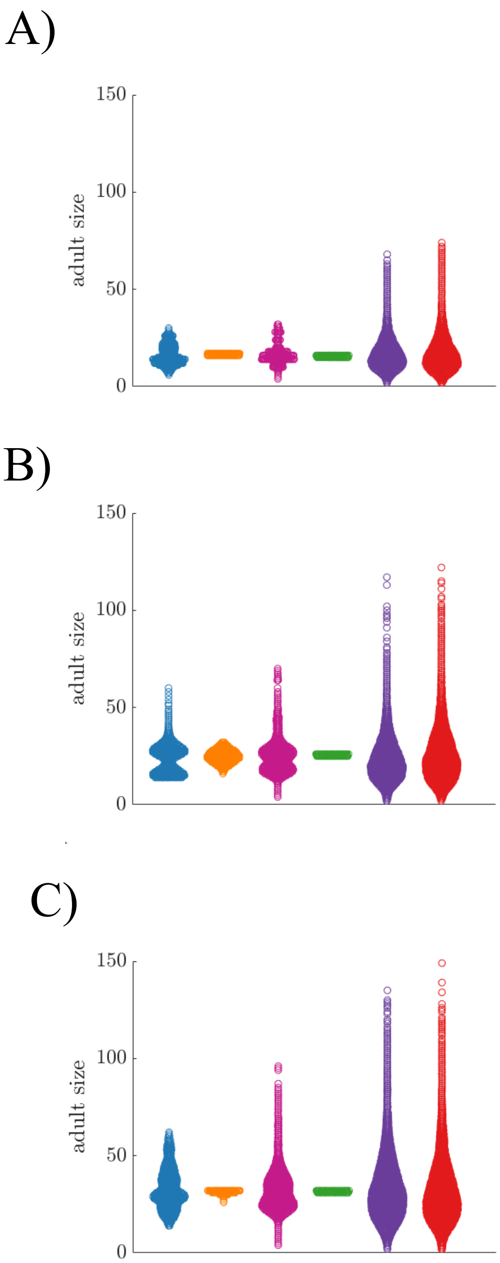


**Fig. S3. Adult size distributions for varying fragmentation thresholds.** A) A swarm chart shows the adult size distribution for values of $\theta$ that results in average adult sizes of 16 cells. B) A swarm chart similar to A) but for average adult sizes of 24 cells. C) A swarm chart similar to A) and B) but for values of $\theta$ that generate average adult sizes of 32 cells, which is the setting that is used in the main paper.

**Reproduction modes for varying $\theta$.** We also analyzed how the fragmentation threshold influences the resulting reproduction modes, see Fig. S4. Overall, we find that the threshold value has little effect on the qualitative fragmentation modes for most information types. Fragmentation based on cell age consistently results in the production of predominantly unicellular propagules. Both connection age and mechanical stress lead primarily to equal binary splits, while the two stochastic information types largely produce unequal binary splits. The only information type for which the reproduction mode shifts with threshold is the diffusing compound. At lower threshold values — corresponding to shorter filaments — it tends to produce equal splits, whereas higher thresholds lead to an increase in unequal binary fragmentation events.

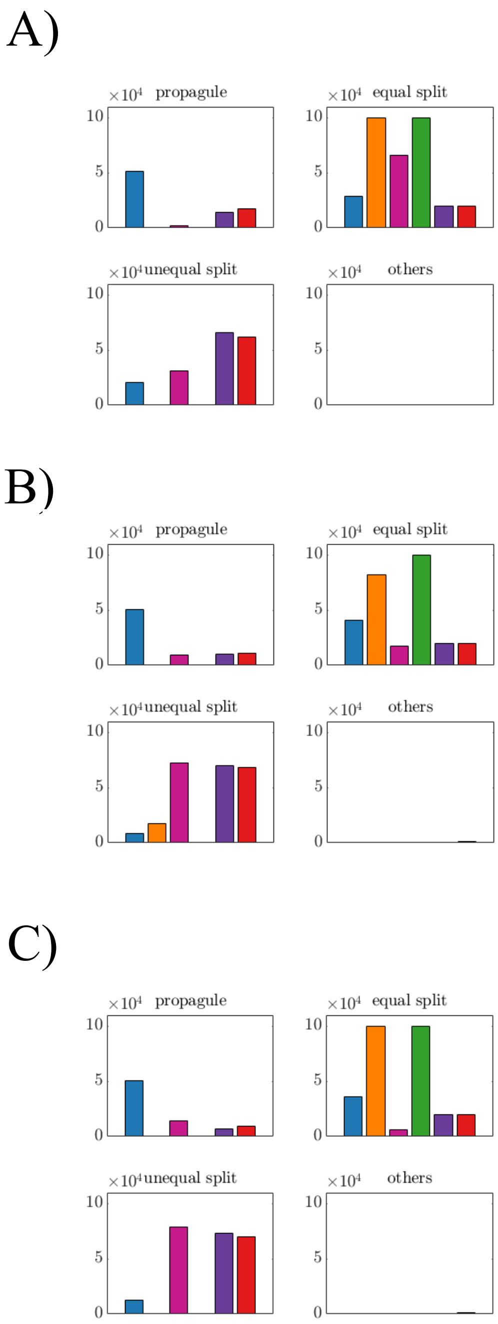


**Fig. S4. Reproduction modes for varying fragmentation thresholds.** A) A bar graph shows the distribution of reproduction modes for values of $\theta$ that results in average adult sizes of 16 cells. B) A bar graph similar to A) but for average adult sizes of 24 cells. C) A bar graph similar to A) and B) but for values of $\theta$ that generate average adult sizes of 32 cells, which is the setting that is used in the main paper.

**Adult size distribution for fragmentation with death.** Our simulations show that including cell death upon fragmentation does not significantly alter the overall adult size distribution. The resulting distributions are presented in Fig. S5. However, all information types experienced a decrease in accuracy, as indicated by a lower proportion of fragmentation events occurring at the optimal adult size of 32.

The most notable change was observed for the cell age information type, where the mean adult size increased by approximately $20\%$, while the standard deviation decreased by $25\%$. Fragmentation with cell death for this this case thus results in larger deviations from the fitness optimum, despite a more consistent fragmentation size — that is, reduced fitness accuracy but increased precision.

For the connection age information type, the mean adult size decreased by $8\%$, and the standard deviation increased three-fold, indicating a notable reduction in fragmentation accuracy. For the remaining information types, fragmentation with cell death led to minor reductions in mean adult size $(1-8\%)$ and slight decreases in standard deviation $(0.5-4\%)$.

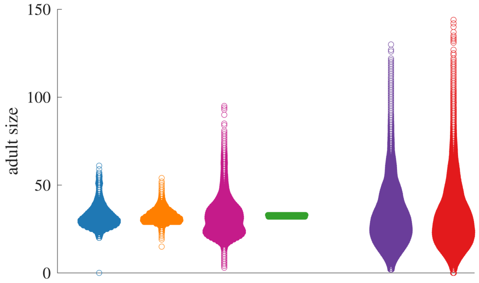


**Fig. S5. Adult size distribution for fragmentation with death.** A swarm chart showing the distribution of adult sizes when fragmentation is a result from cell death. The $\theta$ value is set to generate mean adult size $\approx 32$ in the absence of death.

**Combined information - All combinations.** To explore whether regulation could be improved by combining information, we extended our analysis to pairwise combinations of information types. For each pair, we tested two strategies for combining the information types: an AND function, where both conditions must be met for fragmentation to occur, and an OR function, where fragmentation occurs if either condition is met. While the main paper focused on the combination of cell age and diffusing compound, here we examine all possible pairwise combinations. We find that certain combinations are not possible. In particular, combining stochastic at cell reproduction with age-based information types (cell age, connection age, and diffusing compound) via an AND function results in no fragmentation. This is because the stochastic rule requires newly divided cells, whereas the age-based rules require older cells so it becomes impossible for both conditions to be satisfied simultaneously.

When comparing average adult sizes across combinations (see Fig. S6), we observe that the AND function generally leads to larger-than-target adult sizes, while the OR function tends to produce smaller-than-target sizes. In particular, we find that combining stochastic in time with other information types using an AND function leads to consistently high adult sizes. This is because the deterministic information source has to be fulfilled while at the same time hitting a low-probability event, thus severely delaying fragmentation events.

An analysis of the fitness error (measured via the L2 norm as

$$\log_2\left(\frac{\mathrm{L2\_norm}_{\text{comb}}}{\min(\mathrm{L2\_norm}_{\text{single1}},\mathrm{L2\_norm}_{\text{single2}})}\right),$$

where $\mathrm{L2\_norm}_{\text{comb}}$ is the error for the combined information type and $\min(\mathrm{L2\_norm}_{\text{single1}},\mathrm{L2\_norm}_{\text{single2}})$ is the minimum error obtained when using either information type in isolation; see Fig. S7) shows that combining a well-regulated life cycle (such as diffusing compound or mechanical stress) with another type often increases the error. In contrast, combining two noisy information types can, in some cases, improve regulation and reduce error.

Finally, we examine adult size distributions and reproduction modes for selected threshold pairs $(\theta_1, \theta_2)$ that yield low fitness errors (marked with stars in the L2 norm plots; see Fig. S8). From the adult size distributions, we observe that the AND function can enhance size regulation compared to either information type alone, while the OR function tends to produce an average of the two combined information types. In terms of the reproduction modes, many combinations show an increased frequency of equal binary splits.

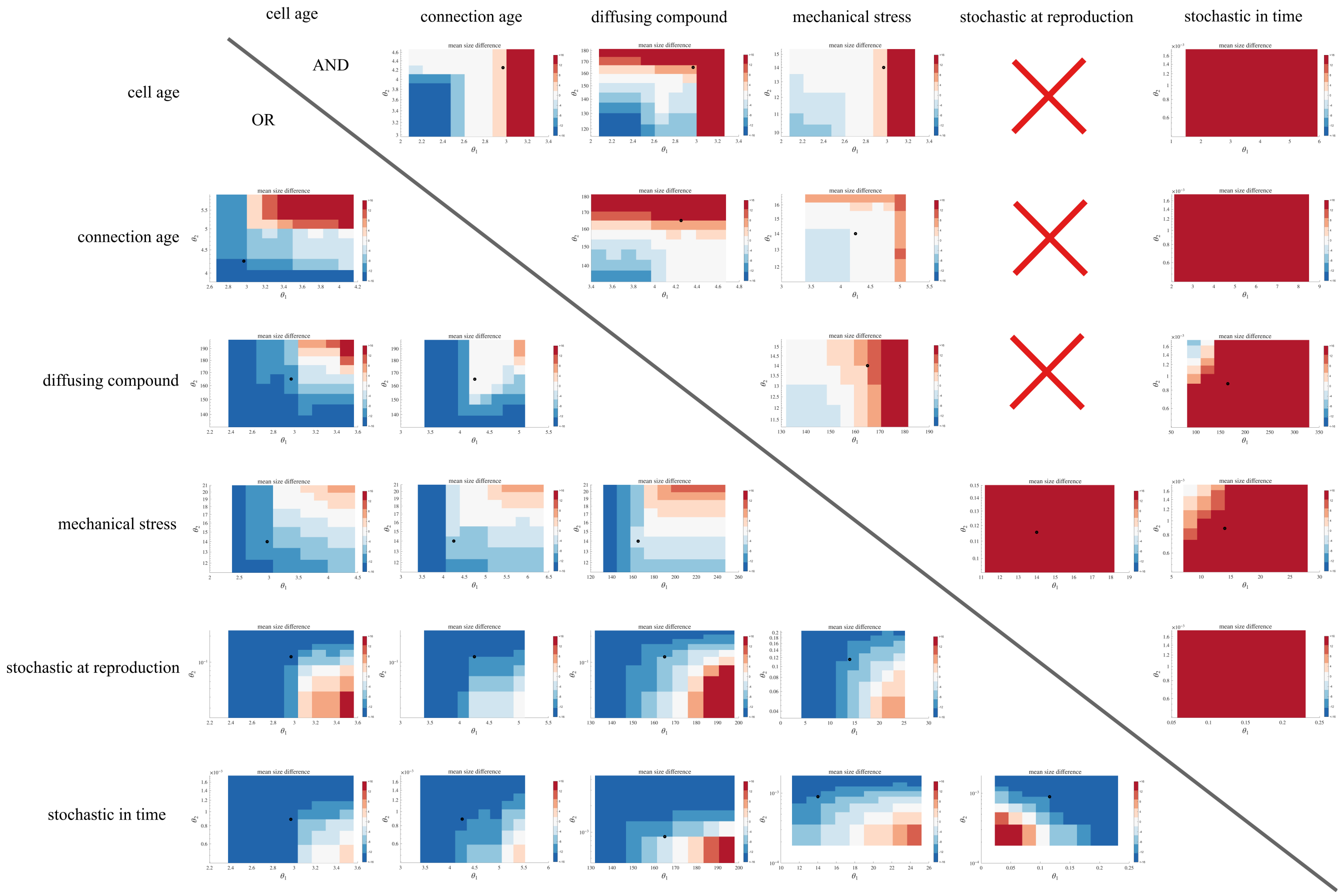


**Fig. S6. Deviation of average adult size from the target size for all pairwise combinations.** Surface plots show the difference between the simulated average adult size and the target size of 32 cells for each $(\theta_1, \theta_2)$ parameter pair. Each subplot corresponds to a specific combination of information types under either the AND or OR function. Red color indicate oversized adults, while blue color reflect undersized adults.

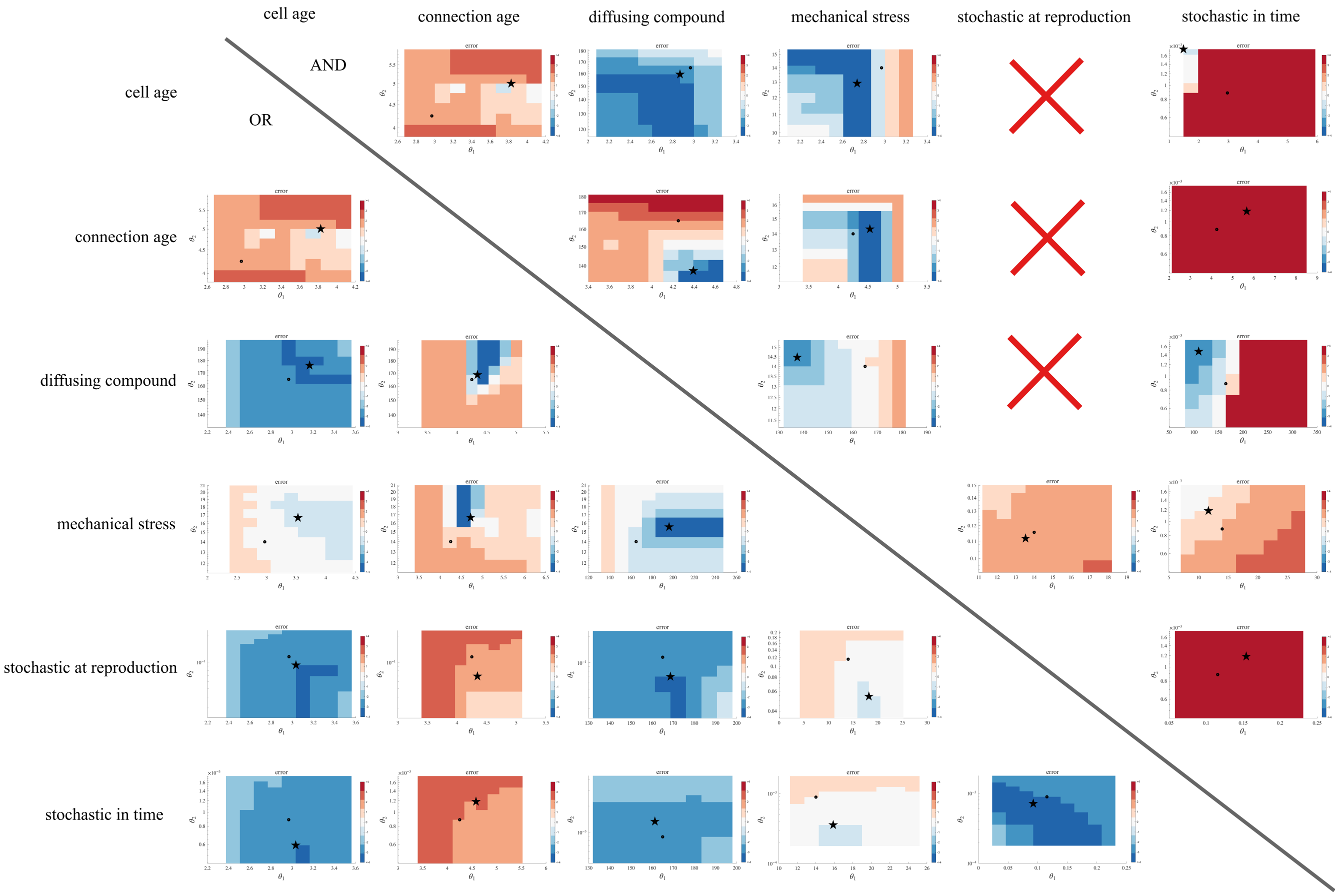


**Fig. S7. L2 norm for all pairwise combinations of information.** Surface plots show the fitness error, calculated via the L2 norm for each $(\theta_1, \theta_2)$ combination. Each subplot corresponds to a specific combination of information types under either the AND or OR function. Red color indicate oversized adults, while blue color reflect undersized adults. Black stars indicate pairs of $(\theta_1, \theta_2)$ with low error, which are selected for further analysis of adult size distributions and fragmentation modes (see Fig. S8).

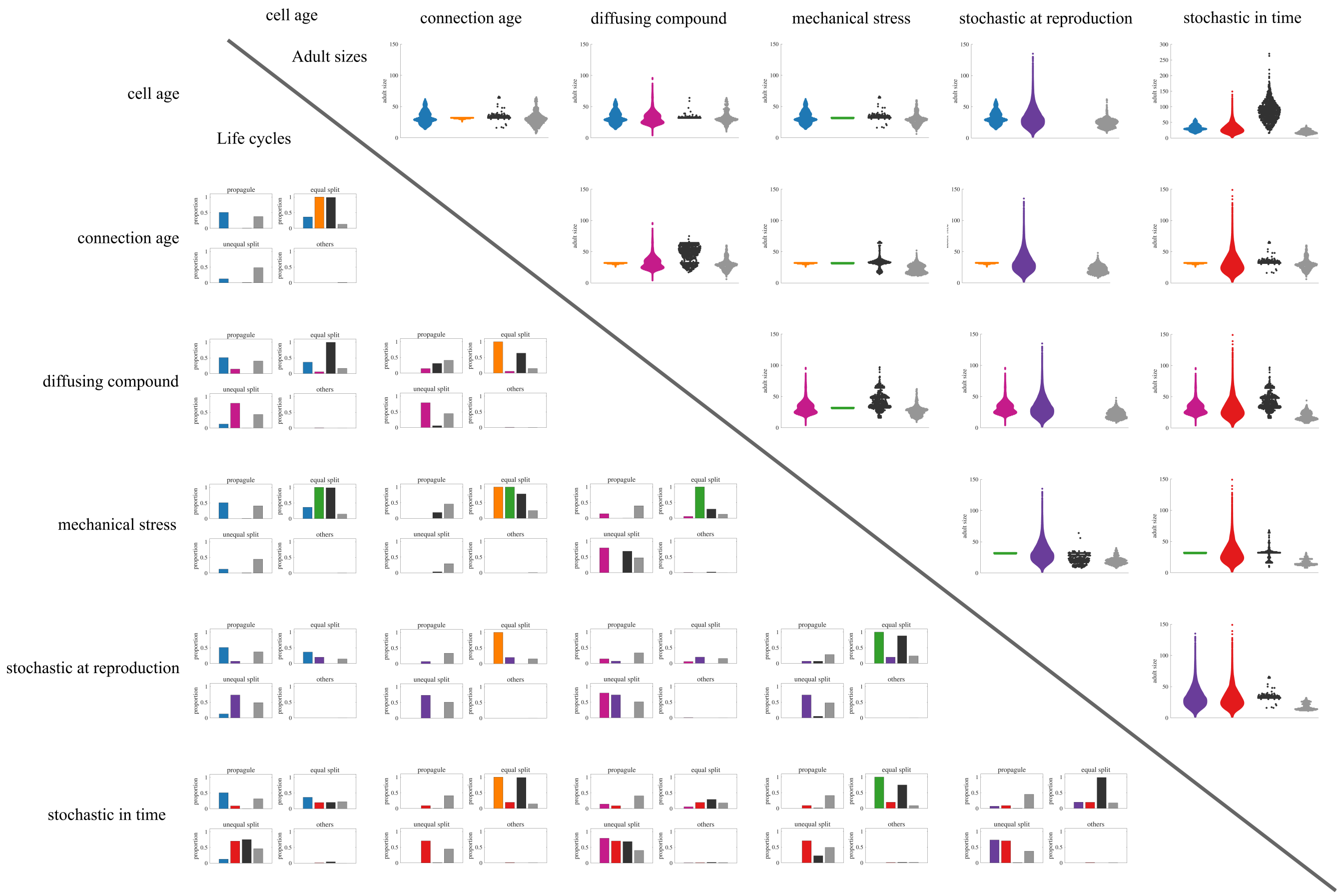


**Fig. S8. Adult size distributions and fragmentation modes for all combinations of information types.** Scatter plots (upper right triangle) show adult size distributions, and bar plots (lower left triangle) show the corresponding reproduction modes for selected $(\theta_1, \theta_2)$ combinations identified in Fig. S7.
